# Age-Related Changes in Neural Networks Supporting Complex Visual and Social Processing in Adolescence

**DOI:** 10.1101/650887

**Authors:** Yulia Lerner, K. Suzanne Scherf, Mikhail Katkov, Uri Hasson, Marlene Behrmann

## Abstract

Despite our differences, there is much about the natural visual world that almost all observers apparently perceive in common. This coherence across observers is evidenced by the finding that, across adults, approximately 30% of the brain is activated in a consistent fashion in response to viewing naturalistic input. The critical question addressed here is how does this consistency emerge and is this pattern of coherence apparent from early in development or does it evolve with time and/or experience? We focused our investigation at a key developmental juncture that might bridge the child and adult patterns, namely, the period of adolescence. We acquired fMRI BOLD data evoked by an 11-minute age-appropriate movie in younger (age 9-14 years) and older adolescents (age 15-19 years) and in adults. Using an intra-subject correlation approach, we characterized the consistency of the neural response within-individual (across two separate runs of the movie), and then, using an inter-subject correlation approach, evaluated the similarity of the response profile within individuals of the same age group and between age-groups. In primary sensory areas (A1+, V1) the response profiles in both groups of adolescents were highly similar to those of the adults, suggesting that these areas are functionally mature at earlier stages of the development. In contrast, some other regions exhibited higher within-age correlations in the adolescent groups than in the adult group. Last, we evaluated the brain responses across the whole cortex and identified the different patterns of maturation as reflected in different inter-subject correlations across the age groups. Together, these findings provide a fine-grained characterization of functional neural development. The approach offers the potential for careful tracking of the development of widespread cortical networks that support the emerging stereotypical responses to naturalistic visual and social stimuli and has important implications for future studies of cortical development.

## Introduction

In spite of our differences, there is much about the surrounding environment that we perceive in common. For example, amongst typical young adults, approximately 30% of the brain exhibits a stereotypical response to viewing the natural world (Hasson et al., 2004; 2010). In other words, a large portion of brain activation is similar or coherent (the term ‘consistent’ is also applicable in this context) in response to the complex inputs we encounter. Importantly, this coherence is not a necessary condition of being a young adult, as individuals with Autism Spectrum Disorder do not exhibit this pattern of neural coherence with each other or with typically developing young adults (Hasson et al., 2009). The question driving the work here is, how do typical adults arrive at these coherent neural responses evident across wide swaths of cortex? Are these patterns of coherence convergent from relatively early in development or do they emerge with time?

### The Development of Neural Coherence

Although there are relatively few studies that measure inter-subject neural consistency across development, there is some evidence that children exhibit stereotypical neural responses when viewing dynamic, naturalistic visual stimuli, but these patterns of coherence differ from those evident in young adults (Cantlon and Li, 2013; Richardson et al., 2018). In one study, children (ages 4-10 years) and young adults watched 20 minutes of Sesame Street video clips. The children show reduced inter-subject consistency (‘neural maturity’ in Cantlon and Li, 2013) in many cortical regions in which adults show the strongest patterns of coherence, including in basic sensory, motor, and association cortices, such as the intraparietal sulcus and Broca’s area. Moreover, the extent to which the coherence deviates from the adult pattern is functionally relevant. Specifically, children who exhibited lower neural coherence relative to adults also exhibited lower math and verbal abilities. There are, however, other regions in which children evince enhanced coherence compared to adults, such as in the superior temporal cortex (Cantlon and Li, 2013). In a similar study, children (ages 3-12 years) and adults watched a short animated film designed to activate regions implicated in understanding other’s minds (i.e., Theory of Mind – TOM regions) and in processing pain. Although the TOM and pain networks were functionally distinct even in the youngest participants, the inter-region correlations increased as a function of age within both networks and the correlations between networks decreased as a function of age (Richardson et al., 2018). These findings suggest that both regions that evince coherence change as a function of age and the extent of that coherence do not necessarily follow a linear trajectory from childhood to adulthood. The critical question then is: what is the trajectory of the changes by which the neural responses of children develop into the adult pattern in which largely coherent neural responses to the natural world are manifest across individuals?

One potential source of variability in the coherence of neural responses amongst children may be related to the ongoing developmental changes that occur in perceptual, cognitive, social, and emotional capacities and the supporting neural circuitry. For example, the perception of real-world objects (Freud and Behrmann, 2017) and the neural basis of size constancy of those objects (Nishimura et al., 2015) continues to evolve throughout childhood. The behavioral (Fry and Hale, 2000) and neural (Scherf et al., 2006) basis of working memory, which could influence the amount information children process as they observe and interpret the natural world, also undergo change. Children’s abilities to process social and emotional information are also changing in impressive ways: the perception of facial expressions improves with age (Motta-Mena and Scherf, 2017, Osterhaus et al., 2016) and the neurophysiology supporting these processes continues to develop at least into early adolescence (Batty and Taylor, 2006). Similarly, higher-order false belief and social understanding (Osterhaus et al., 2016, Devine and Hughes, 2013) undergo age-related improvements as does the functional specialization of networks associated with theory of mind and understanding of pain (Richardson 2018; Richardson et al., 2018). These impressive age-related changes across multiple domains likely contribute to individual variability in the way information is processed, and this variability might result in relatively lower levels of neural coherence across individuals.

As children transition into adolescence, neural organization becomes increasingly adult-like (Luna et al., 2010, Vetter et al., 2014, Moor et al., 2012) and many of the previously described changes in behavior begin to asymptote, including age-related improvements in response inhibition and working memory (see (Luna et al., 2004)). Also, young adolescents (age 12) in later stages of pubertal development, begin to exhibit adult levels of sensitivity to detect emotional expressions (Motta-Mena and Scherf, 2017) and to engage in social reasoning across multiple contexts, like pretense, threat, and irony (see (Vetter et al., 2013)). This convergence in both behavior and neural organization may reflect *increased consistency* in how adolescents process and interpret information. Taken together, these findings suggest one prediction which is that, over the course of adolescence, coherence in neural signals increases between individuals.

However, the opposite prediction may also be feasible as adolescence is marked by two big transitions that arguably influence the perception of the sensory inputs and its neural correlates. First, adolescence begins with the onset of pubertal development (Dahl, 2004), which triggers a cascade of neuroendocrine changes that lead to internal and external physical changes in primary and secondary sexual characteristics and eventual reproductive competence. The earliest aspect of pubertal development, the awakening of the adrenal glands (i.e., adrenarche), begins as early as 6 years of age (Oerter et al., 1990). Gonadarche, which stimulates the release of sex hormones (i.e., estrogens and testosterone), targets the development of the primary sex organs and begins a bit later (typically age 9-10 years; (Susman et al., 2010)). Animal studies indicate large-scale re-organization in structural and functional neural networks that is coordinated with behavior and associated with pubertal development (see (Schulz and Sisk, 2006)) and there is an emerging body of literature implicating the influence of pubertal development on human visuoperceptual behavior (see (Motta-Mena and Scherf, 2017, Picci and Scherf, 2016)), and on functional (Forbes et al., 2011, Spielberg et al., 2014) and structural (Herting and Sowell, 2017) brain development.

Second, adolescence is marked by novel social developmental tasks (SDTs) that facilitate the transition into adult social roles: whereas children are focused on self-mastery and societal propriety and expectations, while still largely dependent on primary caregivers, adolescents are developing increasing autonomy from caregivers as they start to engage in confiding friendships and romantic partnerships with peers. These novel SDTs shape the computation goals of the perceptual system (Scherf et al., 2012, Scherf and Scott, 2012), which influence the kinds of information adolescents perceive, particularly from the social world. Given these two major transitions in adolescence, early adolescence may be a time of increased, rather than reduced, heterogeneity in neural maturity amongst individuals.

### Current study

Because adolescence is a time of substantial psychological, physical, and neural change, we investigated the patterns of variability in stereotypic responses of adolescent brains across the entire cortex as participants watched naturalistic, socially relevant stimuli. To test our predictions about the potential patterns of changes in adolescence, we compared two groups of adolescents to a group of young adults. The younger group of adolescents (ages 9-14 years) were in an age range in which they were likely to be actively undergoing pubertal development, based on age norms in the United States (Susman et al., 2010) and age-related changes in multiple aspects of cognitive, social, and affective development. In contrast, the older adolescents (ages 15-19 years) were in an age range in which individuals were likely to be approaching sexual maturity as well as adult levels of performance on many cognitive tasks. To investigate age-related shifts in neural coherence, in the MRI scanner, blood oxygen level dependent (BOLD) data were collected while participants watched two iterations of the same 11-minute movie clip of children and young adolescents engaging in social exchanges and peer interactions. We compared patterns of within- and between-subject coherence to address three central questions.

First, is there reliability in neural coherence within an individual across the repeated presentations of the movie and does that change with age? There is very little information in the existing literature evaluating test-retest reliability of the neural signal in developmental populations and the extant studies typically have very long intervals (e.g., months to years) between testing points (see (Herting et al., 2017, Cachia et al., 2016)). By assessing the coherence of the entire timeseries across all of cortex across two presentations of the movie and evaluating age group differences in this *intra-subject* coherence, we can explore the reliability and stability of an individual brain across age.

Second, are there differences in neural coherence between the individuals *within* each age group? This allowed us to evaluate how much variability there is among the individuals within each age group and then to determine whether the extent of that within-age variability differed across the three groups. We expected that, across the group of adults, neural coherence would be high especially in sensory regions (primary visual and auditory cortex), superior temporal cortex, and cingulate cortex (Hasson et al., 2004, Hasson et al., 2009). We predicted that, within-group, the younger adolescents would likely exhibit more heterogeneity, and therefore, less neural coherence (and potentially in different regions) given their developmental stage than the older adolescents and adults (see Cantlon & Li, 2013). The patterns of coherence in the older adolescents were harder to predict. Many factors contributing to individual differences may be relatively reduced in the transition from younger to older adolescence, which would be reflected in higher neural coherence. However, older adolescents may still be undergoing significant developmental changes, particularly with respect to SDTs that influence how they make sense of the social visual world, and this could be reflected as reduced neural coherence relative to the group of adults.

Finally, how well does the *between-group* profile of neural coherence in the young adolescents fare relative to that of the older adolescents or adults. This allowed us to determine how different the adolescent groups were in the profile of coherence from each other and from the adult group. Both adolescent groups could differ from the adult coherence profile to an equal extent, but in different ways, or younger adolescents could differ from the adults to a greater extent and in different regions than the older adolescents given the relatively larger developmental differences between the groups.

## Results

The results are presented in three sections, corresponding to the three questions raised above. Figure 1A shows a sequence of images from the movie and the average time course of BOLD activation for each group in the primary visual area (V1).

**Figure 1.**
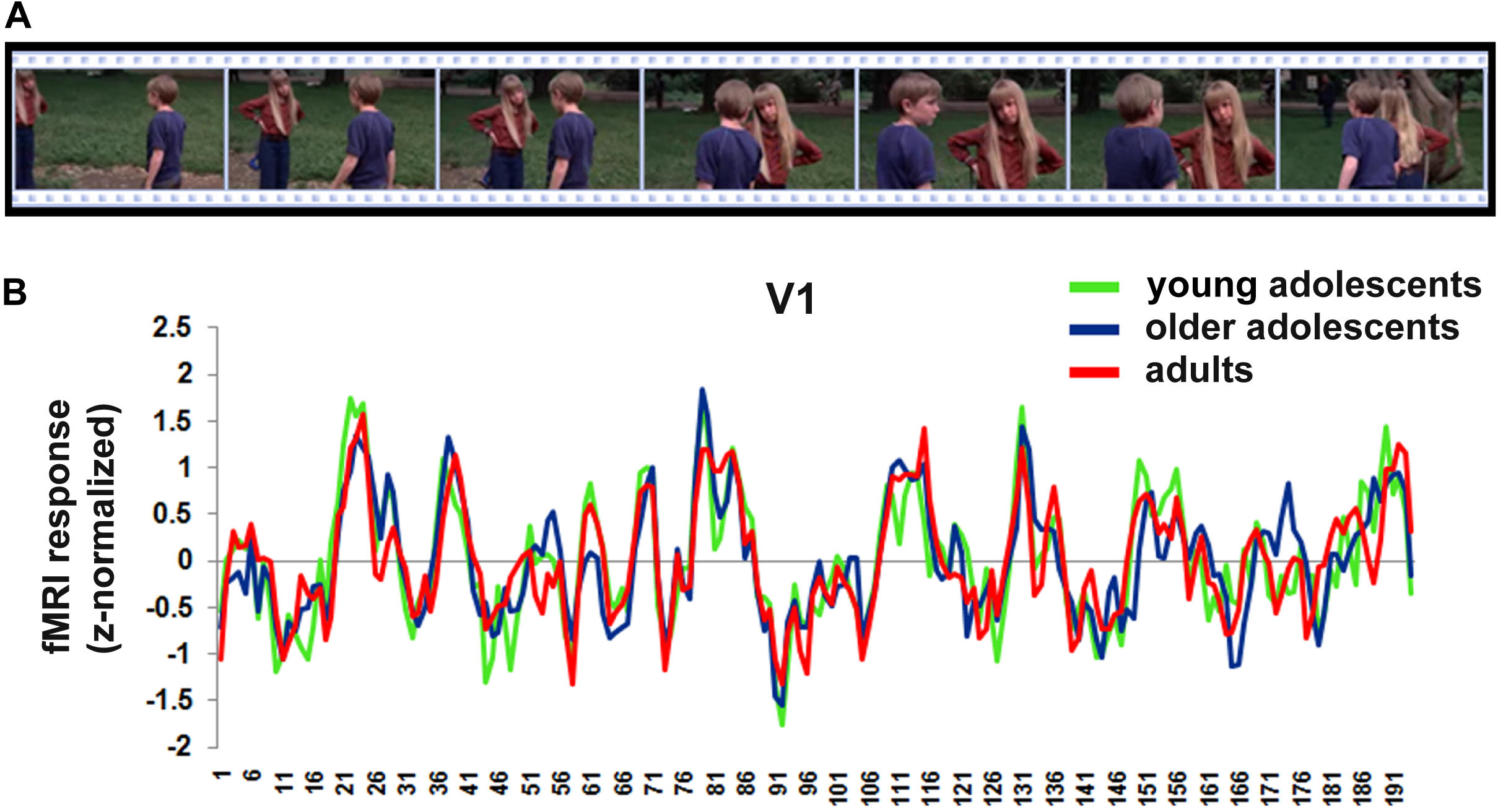
(A) Excerpted sequence from the ‘Escape to Witch Mountain’ movie, which all participants viewed (and heard) in the magnet. (B) The mean z-normalized fMRI response over the course of the movie duration extracted from area VI for all three groups of participants.

### Reliability in Neural Responses within Individuals

First, we assessed whether there is reliability in the neural signals within each individual using *intra-subject correlation* (intra-SC) across the two repeated presentations of the same movie on a voxel-by-voxel basis (see Materials and Methods for details of analysis). Figure 2 (A-C) shows the voxels for each group that exhibited reliable responses across the two versions of the movie. For all three groups, there is reliability in responses of the bilateral ventral temporal cortex, posterior parietal cortex, full length of the middle temporal gyrus, and superior temporal sulcus. To quantify the extent of this reliable response, we computed the percent of brain surface exhibiting reliable responses (i.e. number of voxels exhibiting reliable responses divided by total number of voxels) for each individual and submitted these scores to a one-way ANOVA with group as the fixed factor (Figure 2D). There was no main effect of group, (F(2,24) = 1.6, p = 0.21). Consistent with previous findings in adults (Hasson et al., 2004), young adolescents (M = 25.2%, SD = 7.3), older adolescents (*M* = 26.1%, SD = 8.6), and adults (M = 32.1%, SD = 10.1) all demonstrated coherence in approximately 30% of the cortical surface across repeated presentations of the movie (Figure 2D, left). When we investigated whether age was related to the extent of cortex that exhibited reliable coherent responses within individual in a more continuous way by regressing age in years on the number of voxels exhibiting a reliable response, there was also no significant effect of age, F(1, 25) = 3.0, p = 0.10 (Figure 2D, right). Importantly, although these analyses indicate that an equivalent amount of cortex is reliability coherent in response to the same movie across individuals within each group, they do not evaluate whether *the very same voxels are coherent* across groups.

**Figure 2.**
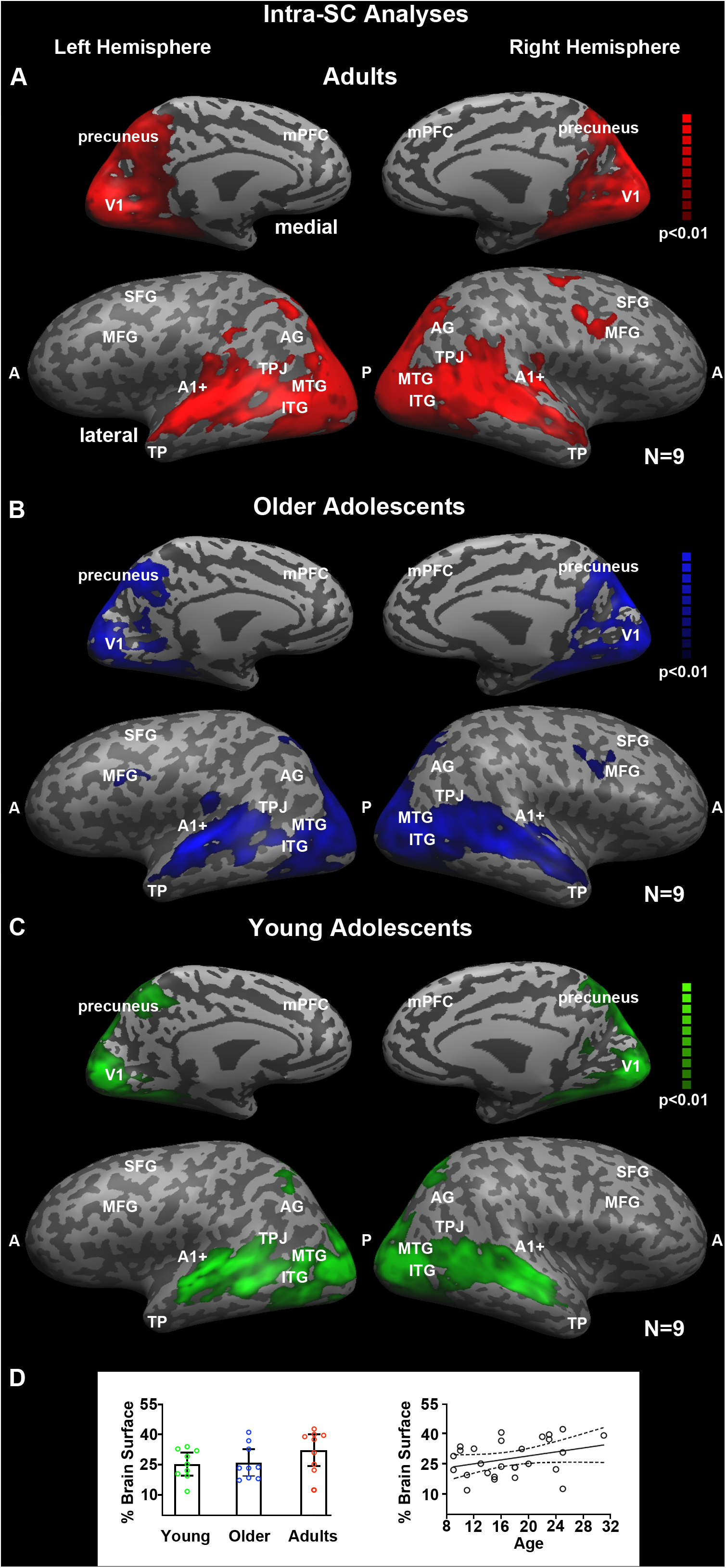
Reliability of responses within participants. These maps illustrate the voxels which exhibit reliable responses across the two presentations of the movie stimulus within individuals as a function of group. The analyses were conducted using intra-subject correlation on a voxelwise basis and corrected using the False Discovery Rate procedure at q < .01. Adults (A), older adolescents (B), and younger adolescents (C) all exhibit reliability in responses bilaterally throughout the ventral visual pathway and full extend of the middle and superior temporal gyri on the lateral surface and medially in the precuneus. (D) The mean percent of brain surface exhibiting reliable responses (i.e. number of voxels exhibiting reliable responses divided by total number of voxels) with 95% confidence interval for each group and each individual as a function of age.

To evaluate this possibility, we compared each pair of intra-subject correlation maps against each other using voxelwise t-tests (see Methods). Figure 3 shows that although there is a good deal of overlap in the sets of voxels that are reliably coherent across within individuals in each group, there are also some important differences. For example, when comparing the adults and the young adolescents (see Figure 3A), it is evident that the adults exhibit more reliability in coherence throughout superior temporal sulcus, angular gyrus, temporo-parietal junction, temporal poles and right superior frontal gyrus. In contrast, the young adolescents exhibit more reliability in coherence in the left intraparietal sulcus. The map of regions where the adults exhibit more reliable coherence than the older adolescents is highly similar to the map of regions where the adults exhibit more reliable coherence than the younger adolescents (see Figure 3B). There are no regions in which the older adolescents show more reliable coherence than the adults. Finally, the older adolescents exhibit more reliable coherence in the precuneus, bilateral middle frontal gyrus, temporal poles, throughout superior temporal sulcus and right temporo-parietal junction compared to younger adolescents (see Figure 3C). However, younger adolescents exhibit more reliable coherence in the left intraparietal sulcus. In sum, these findings indicate that all 3 groups have overlapping but slightly different networks that are reliability activated in response to the same movie across individuals.

**Figure 3.**
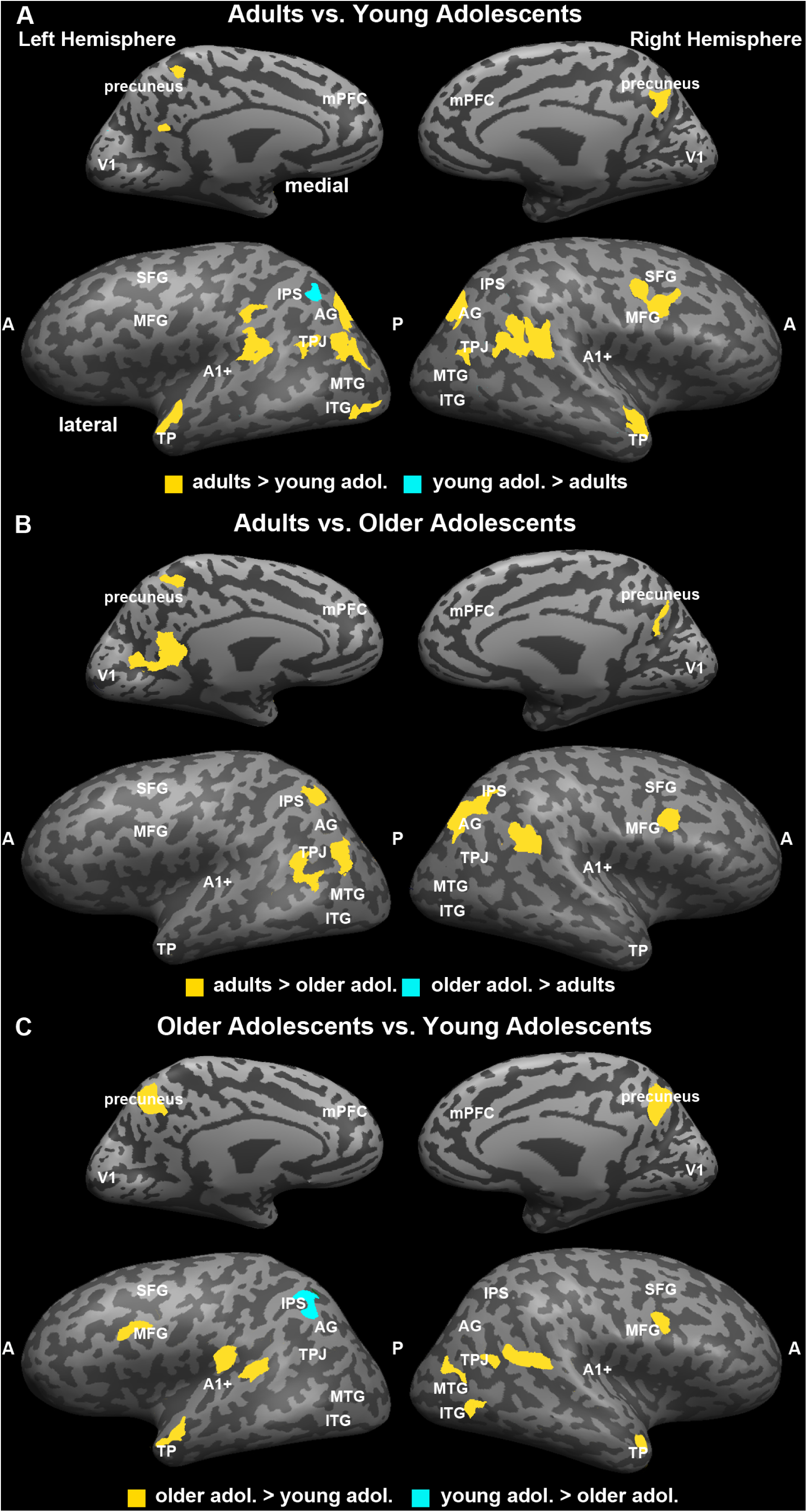
Group comparisons in reliability of neural responses within participants. These maps illustrate pair-wise comparisons of the group reliability maps to evaluate whether the same voxels exhibit reliability in neural responses across the two presentations of the movie stimulus within individuals in each group. In (A) adults and younger adolescents are contrasted, which reveals that adults exhibit more reliability bilaterally in the anterior temporal pole, temporoparietal junction, and in the right superior frontal gyrus on the lateral surface and medially in the precuneus. In (B) adults and older adolescents are contrasted, which reveals that adults exhibit more reliability bilaterally in fewer areas including the temporoparietal junction, and in the right superior frontal gyrus on the lateral surface and medially in the precuneus and posterior cingulate gyrus. Finally, in (C) the older and younger adolescents are contrasted, the differences look much like the contrast between the adults and the younger adolescents.

### Coherence of Neural Responses within Each Age Group

Second, we evaluated the neural coherence of the patterns of activation across individuals *within* each age group in response to the first viewing of the movie. In so doing, we determined how much variability there is among the individuals within each age group and whether that variability differs across the age groups. We used a data-driven approach to investigate the extent to which neural responses across all locations in the cortex are coherent (see Materials and Methods for details of the analysis). This approach has been employed previously to evaluate potential group differences in neural coherence (Hasson et al., 2009, Hasson et al., 2004). Briefly, *inter-subject-correlation* (inter-SC) maps were constructed on a voxel-by-voxel basis by comparing the time course of response for each voxel for a single individual relative to the average time course of the other participants *in the same developmental group*. Finally, for every voxel, the average correlation across individuals of the same group was calculated.

Figure 4 shows the voxel-by-voxel inter-SC maps across the whole cortex both on the lateral and medial surfaces for adults (A), older adolescents (B) and younger adolescents (C). For all three groups there were coherent neural responses among individuals bilaterally in the visual and auditory cortex (see V1 and A1+ labels), throughout the length of the middle and superior temporal gyri, in the inferior parietal lobule and precuneus. Specifically, as evident in the figure, coherent responses were observed (i) in *visually-related* large portions of the posterior cortex, including retinotopic (De Yoe et al., 1996, Sereno et al., 1995, Tootell et al., 1998) and higher-order visual areas (Ishai et al., 1999, Grill-Spector and Weiner, 2014, Hasson et al., 2003), and a region of intraparietal sulcus; (ii) in areas involved in *auditory* processing, and (iii) in the superior temporal gyrus (STG), angular gyrus (AG), supramarginal gyrus, temporo-parietal junction (TPJ), precuneus, inferior occipital gyrus and medial prefrontal cortices.

**Figure 4.**
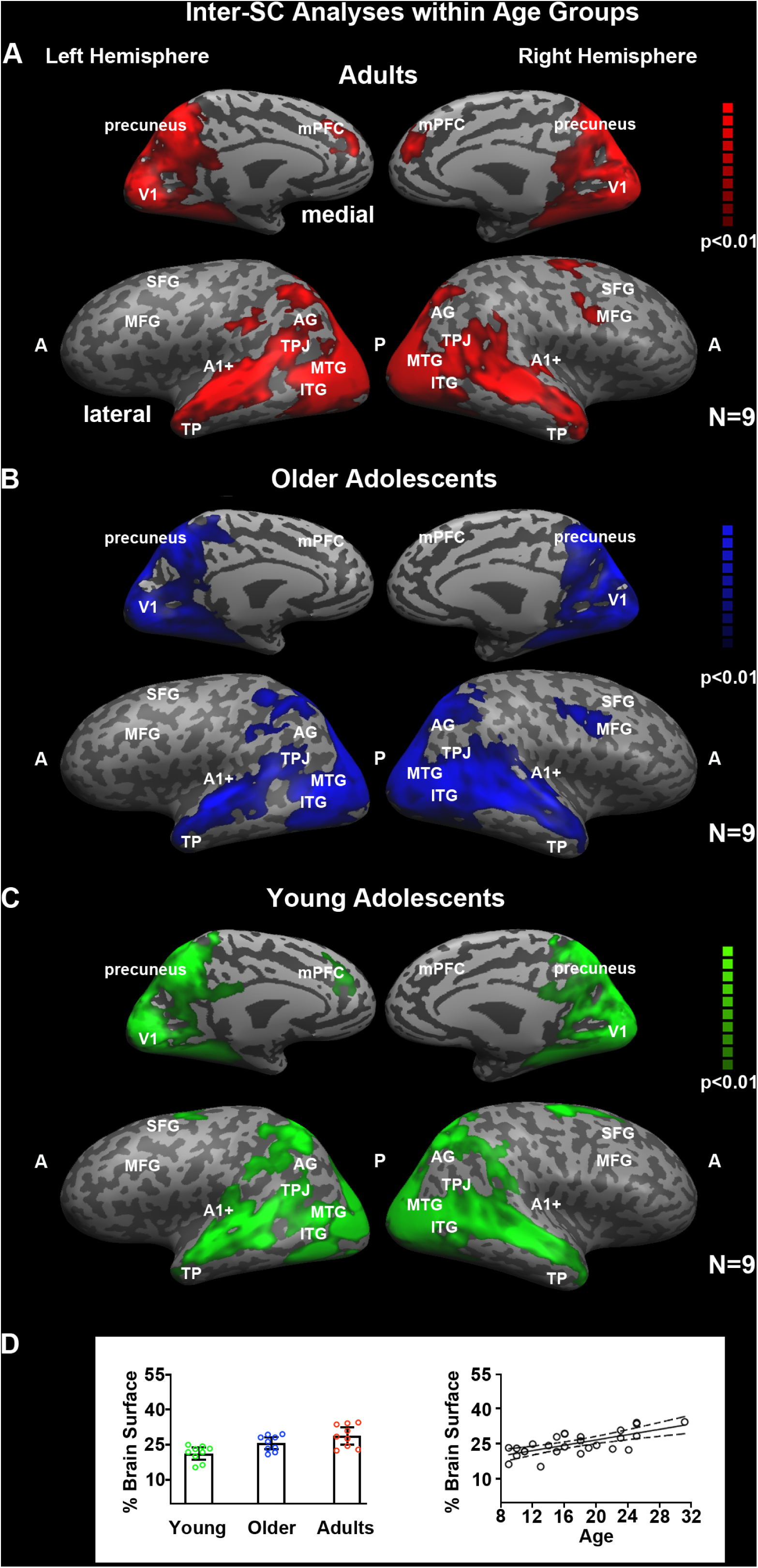
Coherence of neural responses within each age group. These maps illustrate the voxels that exhibit consistency in response profile across individuals within a group in response to viewing the first presentation of the movie stimulus. The analyses were conducted using inter-subject correlation on a voxelwise basis and corrected using the False Discovery Rate procedure at q < .01. Adults (A), older adolescents (B), and younger adolescents (C) all exhibit coherence in responses bilaterally throughout the ventral visual pathway, posterior parietal cortex, and full extent of the middle and superior temporal gyri on the lateral surface, and medially in the precuneus and posterior cingulate gyrus. The adults and young adolescents also exhibit coherence in a dorsal medial prefrontal region. (D) The mean percent of brain surface exhibiting coherence responses (i.e. number of voxels exhibiting coherent responses divided by total number of voxels) with 95% confidence interval for each group and each individual as a function of age. Young adolescents exhibited less coherent response across the cortex than adults and there was a linear trend across the whole age range in increase the amount of cortical tissue that exhibited neural coherence.

A direct comparison of Figure 2 and Figure 4 shows high overlap between the intra-SC and inter-SC maps, indicating that most responses in the posterior, parietal and temporal areas were similar within individual and between the individual participants of the same age group.

To quantify the extent to which coherence in the neural pattern across individuals in the same group varies as a function of the age group, we calculated the mean percentage of voxels that were reliably correlated between individuals within each age group out of total number of voxels (i.e. % brain surface, Figure 4D, left). The overall values for each group: for young adolescents (M = 21.4%, SD = 3.4), older adolescents (M = 25.8%, SD = 3.3) and adults (M = 28.9%, SD = 4.8) (see Figure 4D). We submitted these scores to a one-way ANOVA with group as a fixed factor. There was a significant main effect of group, (F(2,24) = 8.5, p < 0.01). Bonferroni corrected post-hoc tests revealed that only the young adolescents were significantly different as a group than the adults, *p* < .01. We also investigated the relationship between age and percent coherent cortex in a more continuous way using a linear regression with age as the predictor; there was a significant relation, F(1, 25) = 23.1, p < .0001, r^2^ = .48 (see Figure 4D). This indicates that neural coherence between individuals, which may be related to a reduction in individual differences in patterns of neural activation with age.

### Coherence of Neural Responses across Age Groups

To explore differences in the coherence of neural responses across age groups and, specifically, explore, how well the between-group profile of neural coherence in the young adolescents fare relative to that of the older adolescents or adults, we conducted a voxelwise GLM with age group as the fixed factor and subject as a random factor on the z-transformed inter-SC maps. Clusters of voxels (> 25 contiguous voxels) demonstrating a significant main effect of age group (p < 0.05) are shown in Figure 5A (see Table 1 for cluster locations and sizes). The data in Figure 5B illustrate the pattern of the relation between age and the magnitude of inter-subject coherence. However, these data are not independent from the voxel selection process. Therefore, we graph them only to illustrate the nature of the relation between age and change in inter-SCs, not to represent the magnitude of the effect size. We also included V1 and A1+, which were defined anatomically. As a result, the effect of age on the inter-SC in these can be estimated independently.

**Figure 5.**
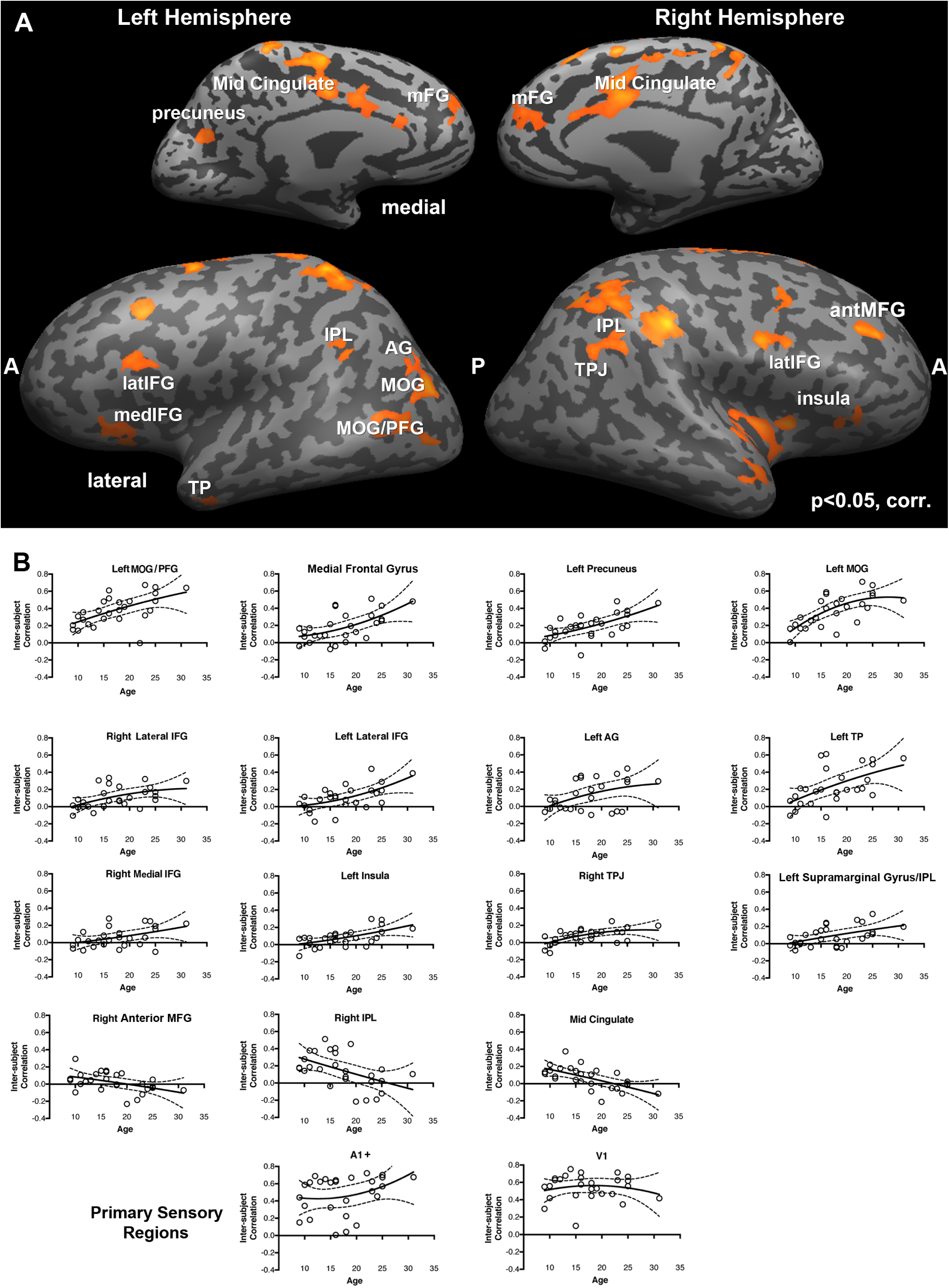
Comparing neural coherence between groups. Group differences in neural coherence, or the consistency in response profile across individuals within a group in response to viewing the first presentation of the movie stimulus. This analysis was conducted by submitting the z-transformed inter-subject correlation maps to a general-linear model with subjects as a random factor and group as a fixed factor. The map was corrected using False Discovery Rate at q < .05. The regions identified as showing an effect of age are shown in (A). To illustrate the direction of the age-related effect in each region, we averaged the inter-subject correlation across the voxels in each region for each participant and plotted the scores as a function of participant age with the best fit line (B). The primary sensory regions were defined separately. A1+ and V1 did not show age-related changes. There were 15 functionally defined regions with age-related effects; 12 of these 15 exhibited age-related increases in neural coherence and 3 exhibited an age-related decline in neural coherence (right IFG, right IPS, middle cingulate).

**Table 1.** Talairach coordinates of the ROIs Exhibiting Main Effect of Age

| ROI | Hem | BA | Talairach Coordinates |  |  | Number of voxels |
| --- | --- | --- | --- | --- | --- | --- |
|  |  |  | x | y | z |  |
| Occipital/Posterior Temporal |  |  |  |  |  |  |
| Middle occipital gyrus (MOG) | L | 19 | -41 | -76 | 12 | 1063 |
| MOG/PFG | L | 37/19 | -50 | -62 | 4 | 780 |
| Parietal |  |  |  |  |  |  |
| Angular gyrus (AG) | L | 39 | -43 | -77 | 25 | 846 |
| Inferior parietal lobule (IPL) | R | 40 | 44 | -32 | 36 | 1217 |
| Supramarginal Gyrus/IPL | L | 40 | -47 | -46 | 33 | 618 |
| Temporoparietal junction (TPJ) | R | 40/39 | 39 | -53 | 31 | 688 |
| Precuneus | L | 31 | -16 | -60 | 17 | 469 |
| Temporal |  |  |  |  |  |  |
| Temporal pole (TP) | L | 21/38 | -53 | 8 | -21 | 172 |
| Frontal |  |  |  |  |  |  |
| Insula | L | 47 | -32 | 23 | -1 | 389 |
| Medial Inferior frontal gyrus (medIFG) | R | 47 | 40 | 27 | 0 | 435 |
| Lateral Inferior frontal gyrus (latIFG) | L | 44 | -46 | 14 | 24 | 437 |
| Lateral Inferior frontal gyrus (latIFG) | R | 44 | 53 | 11 | 33 | 455 |
| Ant. Middle frontal gyrus (MFG) | R | 8 | 35 | 38 | 40 | 494 |
| Medial frontal gyrus (mFG) | Midline | 9 | 0 | 51 | 29 | 859 |
| Cingulate |  |  |  |  |  |  |
| Middle cingulate cortex | Midline | 24 | -2 | 0 | 40 | 1727 |
| Primary Sensory Regions |  |  |  |  |  |  |
| A1+ | L | 21 | -58 | -18 | 1 | 756 |
| A1+ | R | 22 | 55 | -21 | 2 | 326 |
| V1 | R | 18 | 6 | -77 | 1 | 1485 |

Beginning with the independently defined sensory regions, we found no association between age and inter-SC in either V1, F(1, 25) = 0.00, p = 0.99, r^2^ = 0.00, or in A1+, F(1, 25) = 2.14, p = 0.16, r^2^ = 0.07. All three age groups showed fairly large correlations (∼r = 0.5 - 0.6) in these regions. These results indicate that there is comparable inter-subject coherence in the neural signal in these primary sensory regions by early adolescence.

The 15 ROIs identified by the GLM with age as a fixed factor exhibited one of two general patterns of change with age. In 12 out of the 15 ROIs, inter-subject coherence increased with age. These patterns of increasing coherence occur bilaterally in the lateral portion of the inferior frontal gyrus; medially in the prefrontal cortex; in the left middle occipital gyrus, temporal pole, angular gyrus, and precuneus; in the right temporo-parietal junction and in the medial portion of the inferior frontal gyrus. The three regions that show decreasing coherence with age are also distributed and include middle cingulate and right anterior middle frontal gyrus and right inferior parietal lobule. Note, within each region, the association between changes in age and inter-subject coherence are characteristic of the sample and not due to the influence of a few outliers.

## Discussion

This study was designed to investigate age-related changes in the profile of neural coherence as individuals process complex, naturalistic visual input. Moreover, we were specifically interested in whether the mature coherent brain response shared by adults is apparent from early in development or evolves with time and/or experience. In contrast to existing work that investigated neural coherence in young children (Canton & Li, 2015; Richardson et al., 2018), we specifically focused on the presence of age-related changes in neural coherence in younger and older adolescents compared to that observed in adults. As a developmental period, adolescence is accompanied by both behavioral stability and change, in addition to substantial physical change, making it a particularly unique and interesting time to study how neural networks function and potentially re-organize to accommodate these changes within individual and across age. To evaluate neural coherence, we scanned younger (ages 9-14 years) and older (ages 15-19 years) adolescents and young adults using fMRI as they viewed two iterations of a movie clip illustrating social interactions between children and adolescents. We used whole-brain voxelwise analyses to evaluate group differences in patterns of three kinds of neural coherence: first, reliability in the neural signal within individuals as they viewed the two presentations of the movie (intra-subject correlations); second, consistency in the neural signal between individuals as they viewed the movie for the first time (inter-subject correlations); third, consistency in the neural signal between the three groups with a particular focus on the nature of the trajectory across age group in regions of cortex showing significant change across development. Each kind of analysis addressed a separate question about the organization of neural networks across individuals as they process and make sense of complex visual input.

### Reliability of Neural Coherence

There exist limited data on the test-retest reliability of the neural signal in developmental populations, particularly on short time scales (e.g., less than several weeks or months). Here, we assessed the coherence of the entire timeseries for an individual across two presentations of the movie and evaluated potential age group differences in this reliability. We predicted that younger individuals might still be processing the social nuances of the plot through the second viewing of the movie, which could reduce neural coherence across observations compared with the older individuals.

We found that in each of the three age groups, approximately 30% of the cortex exhibited a reliable response across the two presentations of the movie. This finding converges with previous results using this same approach to test the reliability of neural coherence in young adults (Hasson et al., 2004). Importantly, the cortical networks that were reliably used to interpret the movies in each of the age groups were very similar and included the bilateral ventral temporal cortex, posterior parietal cortex, full length of the middle temporal gyrus, and superior temporal sulcus. Interestingly, and different from previous work, all three groups also exhibited reliability in a series of bilateral regions along the precentral sulcus that include posterior portions of the superior and middle frontal gyri.

At the same time, when we directly compared these maps across groups, some age-related differences emerged, particularly in regions implicated in mentalizing and theory of mind processing (Gallagher & Frith, 2003; Saxe & Kanwisher, 2003). Specifically, the adults exhibited more reliable coherence in the left anterior superior temporal sulcus, the right temporal parietal junction and much of the superior temporal sulcus, while the younger adolescents showed more coherence in the middle cingulate gyrus. This pattern of results could indicate a transition in the reliance on the middle cingulate gyrus region over the course of adolescence. In addition, there is also a transition in the reliance on the precuneus, medial prefrontal cortex, and middle frontal gyri with increasing age. Adults exhibited more coherence in these regions than did older adolescents, who in turn, exhibited more coherence in these regions than did younger adolescents. In sum, these findings may reflect an age-related transition in the differential recruitment of regions from the mentalizing network to process information about this movie, which illustrated complex social interactions among peers.

### Individual Differences in Neural Coherence across Age

Next, we evaluated the neural coherence between individuals as they observed the first presentation of the movie. This was computed with the inter-subject correlation and estimated the degree of individual differences in the neural network organization that each group uses to processes the complex visual input. We predicted that adults would evince a similar degree of neural coherence in regions, including primary visual and auditory cortex, superior temporal cortex and cingulate cortex, as has been reported in previous work using this approach (e.g., Hasson et al., 2004, 2009). We also predicted that younger adolescents would exhibit the largest individual differences, given the larger developmental transition that they are undergoing, and, thus, should show the lowest inter-subject coherence throughout most of the neural networks.

Indeed, adults in this study demonstrated comparable neural coherence in sensory areas, like primary auditory and visual cortex, as in previous studies with average inter-subject correlations in A1+ and in V1. Also, we observed extensive inter-subject coherence among all three age groups in the same network of regions in which there was reliability in the neural coherence across iterations of the movie within individuals, including in bilateral ventral temporal cortex and along the lateral surface of the middle and superior temporal gyri. For each of the groups, there were coherent neural responses across individuals in approximately 27% of the cortex, which increased with age. The regions that appear to show the biggest age-related changes in inter-subject neural coherence became evident in the comparison of the inter-SC group maps. The adults show more neural coherence in the mPFC, frontal regions, insular regions, and posterior parietal regions, including the temporo-parietal junction and angular gyrus compared to younger adolescents, who exhibit more coherence in the middle cingulate than adults. Although the cortical networks that the adolescents and adults are using to process and interpret the complex visual and social information in the movie is largely overlapping, the important differences include regions in the mentalizing network (i.e., mPFC, middle cingulate, TPJ, angular gyrus/posterior superior temporal sulcus). These findings converge with those from the analyses of the reliability of the neural signal to suggest that this movie invokes networks implicated in mentalizing in all three age groups and that there is differential weighting and/or recruitment of the specific regions in these networks that changes with age.

Finally, we compared the neural coherence between individuals in every voxel of the brain. This allowed us to generate a map with clusters of voxels that represented age-related changes in the magnitude of the inter-subject correlation. We predicted that there might be nonlinear differences in the way each of the adolescent groups differ from the adult group given their different developmental profiles. The majority of the clusters revealed a pattern of increasing neural coherence with age. These clusters were distributed throughout the brain and included many of the same regions that were identified in the previous neural coherence analyses, such as mPFC, precuneus, right TPJ, left temporal pole, and right insula (pars triangularis). In contrast, there were three regions in which the neural coherence decreased with age, including in the right anterior middle frontal gyrus, middle cingulate, and right inferior parietal lobule. Again, these are regions that have been consistently implicated in mentalizing and theory of mind processing in both adults and children (for review see Mahy et al., 2014).

### Age-Related Changes in Neural Networks Supporting Complex Visual and Social Processing in Adolescence

In sum, across all three metrics of neural coherence, our results converge to reveal two central findings about the neural networks adolescents and adults use to process complex visual and social information. First, the cortical networks invoked as adolescents and adults process complex visual and social input are largely similar and include higher-order object processing regions in the ventral visual pathway and regions implicated in processing social interactions that are distributed throughout the brain (see Redcay & Warnell, 2018). Second, age-related differences in these networks are largely found in the extent to which individuals rely on particular regions within these networks. Specifically, younger adolescents rely more on the middle cingulate gyrus and less on the mPFC, precuneus, TPJ, insula, and frontal regions than do adults when processing and interpreting these complex visual scenes involving social interactions with peers.

Given the series of findings we have reported, the implications for the two proposed predictions become clear. On the one hand, we have obtained evidence suggesting that the coherence profile of the brains of individuals in adolescence seems well suited to bridge between childhood to adulthood: in multiple cortical regions, the younger adolescents’ brain responses showed less consistency to each other than was true of the older adolescents who, in turn, showed less consistency in neural responses than adults. Even close scrutiny of those clusters where the effect of age is most prominent reveals an incremental development of coherence across age. These results are consistent with the findings of many studies which report an increasing progression in cognitive, social, and emotional capacities and the supporting neural correlates. The findings are not easily accounted for by a framework in which heterogeneity of neural responses dominates the coherence amongst brains of adolescence and, subsequently, share little commonality with the coherence profile among adult brains. Although adolescence is a time of large-scale re-organization in neural networks as a result of pubertal development and challenging social developmental tasks (Blakemore et al., 2010), the results of the present investigation favors an interpretation that young adolescents use overlapping but slightly different networks when making sense of the movies. This is evident in a couple of findings: the majority of cortex is not different (all grey regions) in inter-SC revealing that the networks are not that different. However, closer scrutiny reveals that the middle cingulate and right IPL have higher intra-SC in the younger adolescent group across the two viewings of the movie and both show decreasing inter-SC with age during run 1, that the youngest group relies on these regions more than the two older groups do in order to interpret visual stimuli. This suggests that these regions become less important with age. In contrast, the two older groups have more intra-SC in left angular gyrus and posterior left inferotemporal gyrus from across the two viewings of the movie, and the inter-SC in these regions increase with age – indicating that these regions become more integrated into the network with age. It is worth noting, however, that the finding reveal both positively and negatively linear functions of brain coherence across age, with the latter situated closer to frontal cortex and regions associated with executive function and working memory.

Together, these findings therefore offer a more nuanced view of development and suggest that maturation is not an all-or-none phenomenon nor is it the case that maturation is uniform and that all brain regions obey the same functional profile. It is well-established that higher-order association cortices mature after lower-order somatosensory and visual cortices (Gogtay et al., 2004) and that the structural co-variance of networks (in which cortical thickness in one region influences the thickness of structurally and functionally connected regions) is coordinated or synchronized over the course of development (Alexander-Bloch et al., 2013). However, not all facets of brain development are in lockstep, unfolding in a progressive sequence. For example, individual sulcal patterns of anterior cingulate cortex (ACC) are fixed from childhood to adulthood, even though quantitative anatomical ACC metrics may be changing dramatically (Cachia et al., 2016).

More recently, structural differences across age have been explored by computing the similarity in the trajectory of cortical thickness within individual and across age between any two cortical regions, a profile of maturational coupling (Khundrakpam et al., 2019). Using this approach, the authors report that changes in brain structure occur in a coordinated fashion over time, that these changes are well aligned with functional connectivity (as in the default mode network) and that this individual-based structural covariance approach offers the possibility of tracking variability and may provide a mechanistic explanation in cases with neural (and neurodevelopment) disorders. With the advent of innovative analytic approaches, whole-brain data can be relatively easily explored and analyses of individual and group similarity or variability better elucidated. Together, the findings reported here characterize the fine-grained developmental trajectories of regions of cortex engaged by naturalistic stimuli (as far as is possible in the bore of the magnet) and put a spotlight on the specific challenges of cortical development as the adolescent brain approximates the more mature, stable brain profile of adulthood.

## Materials and Methods

### Participants

Participants included 9 young adolescents (age: 9-14 years; *M* = 11.0, *SD* = 1.7; 7 males), 9 older adolescents (age: 15-19 years; *M* = 16.5, *SD* = 1.4; 3 males), and 9 adults (age: 20-31 years; *M* = 24.0, *SD* = 3.0; 5 males). An additional 3 young adolescents, 3 older adolescents, and 2 adults were excluded from the analyses due to excessive head motion (> 2.8mm), technical problems during the scan (e.g. image distortion, no sound) or request to stop the scan prior the end. One additional adult, randomly chosen, was excluded to equalize group size (inclusion of this subject did not change the results). All participants were healthy, with no history of neurological or psychiatric disorders in themselves or in their first-degree relatives, had normal or corrected vision, and were right-handed, native English speakers. Prior to participating in the study, participants and/or their guardians provided assent and/or written consent. The protocol was approved by the University of Pittsburgh and Carnegie Mellon University Internal Review Boards.

### Stimuli

Movie stimuli were created by extracting two clips from a G-rated movie, *Escape to Witch Mountain* (1975, Walt Disney Productions). We specifically chose a movie with a plot and characters that were understandable by the youngest age group in order to minimize the likelihood that potential differences in patterns of neural activation were related to differences in comprehension of the movie (see also Alexander et al., 2017). We also selected an older movie (i.e., produced before any of the participants was born) that we thought was likely to be novel to the majority of participants to minimize interactions between age group and familiarity that might impact patterns of neural activation.

### Procedure

Immediately prior to the scanning session, all participants were trained in a mock scanner for 15 - 30 minutes. Each participant practiced lying still while watching a movie inside the mock scanner with simulated scanner noises. Participants were instructed to use relaxation breathing, encouraged to use mental imagery (e.g., lying in own bed watching a movie), and provided with feedback about when they moved during the simulation. This simulation procedure acclimates participants to the scanner environment, minimizes motion artifacts, and reduces anxiety in both children and adults (Scherf et al., 2014). During this simulation session, participants watched the first 10 minutes of the movie to promote an understanding of the plot prior to observing the experimental excerpt. The next 11.5 minutes of this movie were selected as the stimulus for the experimental scanning paradigm.

Participants were scanned at the Brain Imaging Research Center in Pittsburgh on a Siemens 3T Allegra Scanner, which was equipped with a quadrature birdcage head coil. During the scanning session, the movie stimuli were displayed using QuickTime on a rear-projection screen located inside the MR scanner. Participants wore MRI-compatible headphones and passively viewed stimuli in three functional runs, including two identical iterations (Run 1 and Run 2) of the Movie task (see Figure 1) (and a localizer scan which is not being analyzed for this study). To ensure that the participants were watching the movie throughout, we monitored their eye movements in the scanner using an infrared video camera equipped with custom built MRI telephoto lens (Applied Sciences Laboratories, Model504LRO). The total scanning time was approximately 45 minutes. Following the scan, outside the magnet, participants completed a set of questions about the characters and plot of the movie to verify that they understood the narrative. All participants exhibited good comprehension of the plot and, therefore, we included the data from all of these participants in the analysis.

### MRI Scanning

Functional echo-planar imaging (EPI) images were acquired in 35 AC-PC aligned slices that covered most of the brain (TR = 3000 ms; TE = 35 ms; 64 × 64, 3 mm slice thickness, 3.2 × 3.2 mm in-plane resolution). High-resolution anatomical images were also acquired during the same scanning session using a 3D-MPRAGE pulse sequence with 192 T1-weighted, straight sagittal slices (1 mm thickness).

## Image Processing and Data Analysis

### Preprocessing

Data analyses were performed using BrainVoyager QX (Brain Innovation, Maastricht, Netherlands) together with in-house software written in MATLAB. Preprocessing of the functional data included 3D-motion correction, filtering out of low frequencies (e.g. slow drift) up to 10 cycles per experiment, linear trend removal, high-pass filtering (cut-off: 0.01 Hz), and spatial smoothing with a Gaussian filter (6 mm full-width at half-maximum value). The first 25 and last 8 TRs in each run were removed to eliminate pre-processing artifacts and to allow the hemodynamic responses to reach a steady state. The time-series data for each run of the task for each participant were then spatially normalized into Talairach space, an approach that has been validated in a previous developmental study (Burgund et al., 2002), and projected onto a reconstructed cortical surface from the high-resolution 3D anatomical images.

Only participants who exhibited motion of less than one voxel (3 mm) in all six directions (i.e., no spikes in motion greater than 2.8 mm in any direction on any image) for both runs of the movie were included in the fMRI analyses. In addition, we evaluated whether the mean motion in each of the six directions varied as a function of age group. Mean motion did not exceed 0.5 mm (approximately 1/5 voxel) in any individual in any direction. Maximum motion aggregated across all participants in all directions was 1.7 mm. Separate one-way ANOVAs on each of the motion vectors revealed that there were no group differences in motion in any direction for either functional run of the movie (all p > 0.1). However, separate linear regressions using age to predict motion parameters revealed that age was negatively associated with rotation along the Z axis during the first functional run of the movie, F(1, 26) = 4.53, p = 0.043.

### Reliability of Neural Responses within Individuals Across Movie Presentations

Following our previous work (Hasson et al., 2008), intra-SC was computed in each voxel over the entire cortex for each participant as follows: 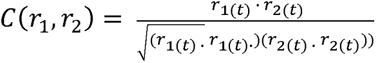 where r_1_(t) and r_2_(t) are the response time courses of a voxel each of the presentations of the movie.

### Coherence of Neural Responses between Individuals within Each Age Group

The inter-subject correlation (inter-SCs) maps were also constructed on a voxelwise basis within each age group. In each subject, the Pearson product-moment correlation was computed between the raw voxel time course and the average voxel time course from the remaining participants so that each participant was excluded from the average they were being compared to (see Hasson et al., 2004, 2008, 2010). Next, the inter-SC was generated in each age group by computing an average correlation across participants for each voxel. The analysis revealed systematically stronger correlations in Run 1 than in Run 2 for all three age groups. As a result, the inter-SC data from Run 1 was used to quantify and compare findings across the groups.

### Coherence between Age Groups as Compared to Adults (Neural Maturity)

To evaluate how each adolescent group differed from the adult group as a measure of neural maturity as in previous work (Cantlon & Li, 2015; Richardson, et al., 2018), inter-SC maps were computed for each individual in each adolescent group on a voxelwise basis relative to the adult time course in each voxel. Next, the average correlation was calculated across participants in every voxel separately within each age group. In addition, because we were interested in age-related changes from the young to the older adolescents, we performed the same correlation for the younger versus older adolescents in each voxel.

### Assessing Significance of Maps

The statistical significance of the correlation map of each individual was assessed via a phase-randomization procedure (for recent review, see Poldrack 2017). Phase-randomization was performed by applying a Fast Fourier Transform to the signal, randomizing the phase of each Fourier component, and then inverting the Fourier transformation. Thus, the power spectrum was preserved but the correlation between any pair of such phase-randomized time courses had an expected value of 0. Phase-randomized time courses were generated for every measured fMRI time course from every voxel in each participant. A correlation value was then computed (for each kind of analysis described above) for every voxel. This permutation process was repeated 5000 times to generate a null distribution of the correlation values, separately for each voxel. Statistical significance was assessed by comparing the empirical derived correlation values from each analysis with these null distributions. The false discovery rate (FDR) was employed to correct for multiple comparisons (Benjamini and Hochberg, 1995, Benjamini et al., 2001, Genovese et al., 2002).

Group differences between correlation maps were evaluated by computing voxel-wise independent-samples t-tests between each pair of groups (e.g., adults vs. older adolescents, adults vs younger adolescents). The voxel-wised t-test was done by comparing the Fisher-transformed correlation values computed in each voxel of participants from different groups.

Finally, to evaluate how neural coherence changes as a function of age, each individual inter-SC map was submitted to a Fisher Z-transformation so that they could be submitted to a single, voxelwise GLM with age as a fixed factor. Voxels that were significantly active at a *p* > .05 and that existed in a cluster of 25 or more contiguous voxels were identified. This analysis revealed that age was significantly related to the magnitude of the inter-SC in each clusters of voxels. However, it did not reveal what the specific pattern of change across age was in any cluster. To illustrate these patterns, we computed the average inter-SC for each participant across the set of voxels from each significant cluster and plotted the scores against age with the best fit quadratic function and 95% confidence intervals.

We also identified two primary sensory regions anatomically, including early auditory cortex (A1+) and early visual cortex (V1), to examine group differences in the inter-SC. The early auditory cortex cluster (A1+) was defined as the set of voxels that correlated most highly with the stimulus audio envelope. The early visual cortex (V1) was defined anatomically. Clusters for these primary cortices (A1+ and V1) were included as these are strongly correlated in adult brains and therefore provide a benchmark for a high degree of consistency among observers (Hasson et al., 2004). To compute the correlation between the average BOLD signals and the audio envelope, we bandpass filtered the audio signal from the movie between 4 and 4000 Hz, extracted the envelope of the signal using a Hilbert transform, and then down sampled the envelope to the sampling rate of the BOLD signal using an anti-aliasing lowpass finite impulse response filter.

## Acknowledgements

This research was supported by a grant to MB from the National Institutes of Health (R01EY027018-NEI-BEHRMANN).

